# Streamlining differential exon and 3’ UTR usage with diffUTR

**DOI:** 10.1101/2021.02.12.430963

**Authors:** Stefan Gerber, Gerhard Schratt, Pierre-Luc Germain

**Author notes:** Correspondence to Pierre-Luc Germain.

## Abstract

**Background:** Despite the importance of alternative poly-adenylation and 3’ UTR length for a variety of biological phenomena, there are limited means of detecting UTR changes from standard transcriptomic data.

**Results:** We present the *diffUTR* Bioconductor package which streamlines and improves upon differential exon usage (DEU) analyses, and leverages existing DEU tools and alternative polyadenylation site databases to enable differential 3’ UTR usage analysis. We demonstrate the *diffUTR* features and show that it is more flexible and more accurate than state-of-the-art alternatives, both in simulations and in real data.

**Conclusions:** *diffUTR* enables differential 3’ UTR analysis and more generally facilitates DEU and the exploration of their results.

## Background

Coding sequences in eukaryotic mRNAs are generally flanked by transcribed but untranslated regions (UTRs) which can impact RNA stability, translation, and localization^[1]^. In particular, the length of 3’ UTRs often varies even within a given gene due to the use of different poly-adenylation (polyA) sites^[2]^, leading especially to the inclusion or not of regulatory elements such as binding sites for microRNAs (miRNAs) or RNA-binding proteins^[3]^. Alternative poly-adenylation (APA) is highly prevalent in mammals^[4]^ and has been shown to be important to a variety of biological phenomena^[5,6,7,8]^.

A number of methods for 3’ end sequencing have been developed with the goal to map APA sites^[9,10,11,12,13,4,14]^, leading to the development of atlases such as *PolyASite*^[15]^ or *PolyA_DB*^[16]^. As such methods are only marginally used, however, it would be beneficial to leverage the widespread availability of traditional RNA-seq for the purpose of identifying changes in 3’ UTR usage. A chief difficulty here is that most UTR variants are not catalogued in standard transcript annotations, limiting the utility of standard transcript-level quantification based on reference transcripts, such as *salmon*^[17]^. Nevertheless, a number of methods have been developed to this purpose. Methods like *DaPars*^[18]^ and *APAtrap*^[19]^ try to infer new polyA sites from read coverage changes from RNA-seq experiments, however the depletion of RNAseq coverage at the 3’ end of transcripts makes the precise inference of polyA sites challenging^[20]^. Other tools like *QAPA*^[8]^ and *APAlyzer*^[21]^ use already available polyA site databases but only compare the usage of the most proximal polyA sites to distal ones in a pairwise fashion and fail to grasp the full complexity of dynamic APA when there are three or more polyA sites, which is the case for approximately half of mammalian transcripts^[4]^. Furthermore they do not make use of the already proven statistical frameworks to analyse different exon usage (DEU) from count data^[22,23,24,25]^. These tools take into account the inherent properties of read count distributions and are arguably more appropriate to analyse differences in relative polyA site usage, which is conceptually highly similar to DEU. We therefore developed *diffUTR,* which streamlines and improves upon well established DEU tools, and leverages them, along with polyA site databases, to infer alternative 3’ UTR usage across conditions.

## Results

### Streamlining differential bin/exon usage analysis

Popular bin-based DEU methods are provided by the *limma*^[25,24]^, *edgeR*^[23]^ and *DEXSeq*^[22]^ packages. However, their usage is not straightforward for non-experienced users, and their results often difficult to interpret. We therefore developed a simple workflow (Figure 1A), usable with any of the three methods but standardizing inputs and outputs. In particular, bin annotation and quantification, as well as different usage results, are all stored in a `RangedSummarizedExperiment`^[26]^, which facilitates data storage and exploration, and enables advanced plotting functions irrespective of the underlying method. *diffUTR* is flexible in its application, and supports the use of strand information if available.

**Figure 1:**
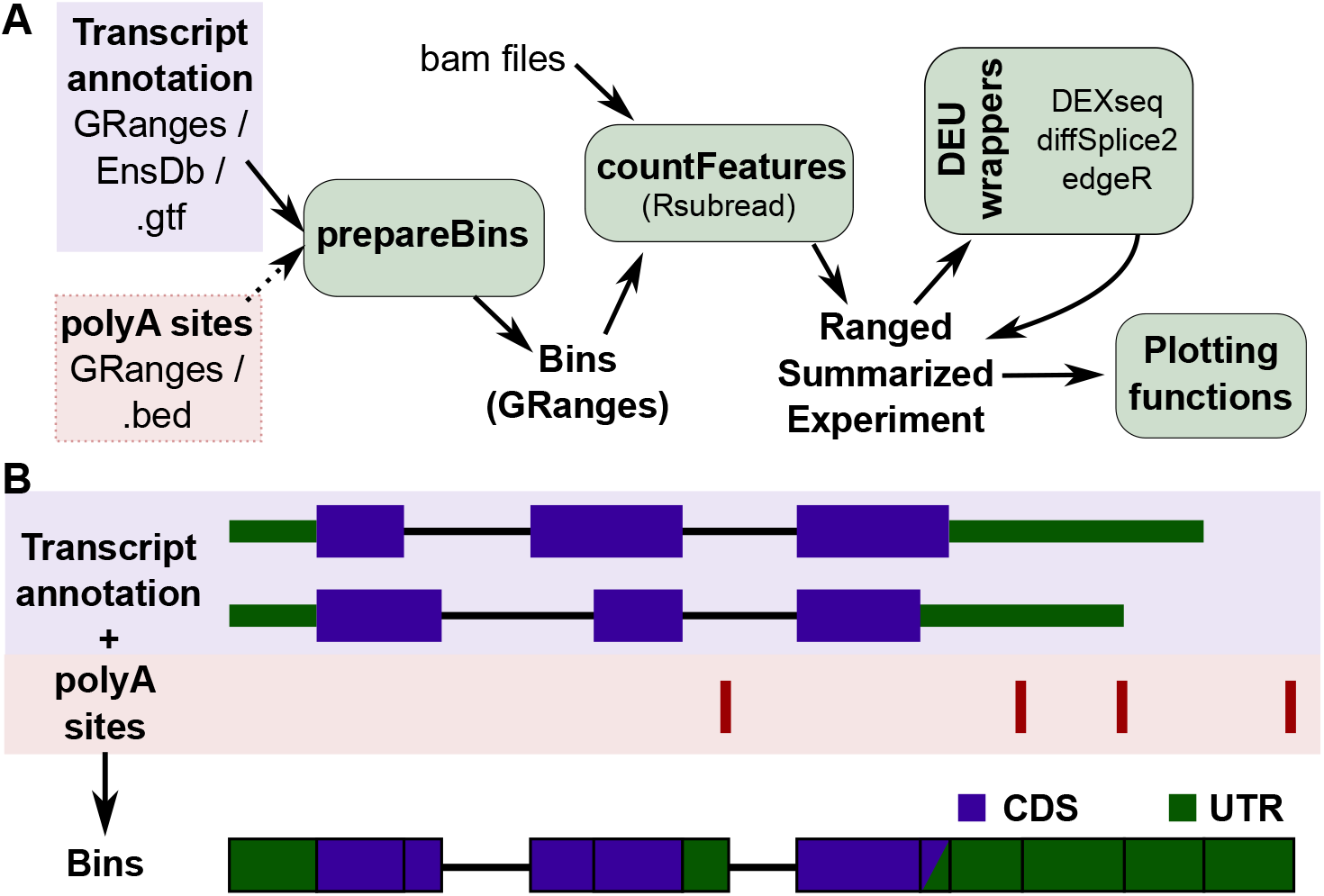
Overview. **A:** *diffUTR* workflow. Bins are prepared from various types of gene annotations as well as, optionally, additional APA-driven segmentation and extension, then read counts within bins as well as bin information are stored in a standardized `RangedSummarizedExperiment`, which can then be used as an input for any of the three DEU methods, producing again a standardized output that can be used with the package’s plotting functions. **B:** Schematic of bin preparation. APA sites are used to further segment and extend disjoined gene bins.

### Improvement to diffSplice

*diffUTR* also implements an improved version of *limma’s* diffSplice method which does not assume constant residual variance across bins of the same gene (see diffSplice2). To test the effect of these modifications in a standard DEU setting, we ran both versions (as well as the other two DEU methods) on simulated data from a previous DEU benchmark^[27]^. The precision and recall results (Figure 2A) confirmed the previously observed superiority of *DEXSeq* and, more generally, the imperfect false discovery rate (FDR) control. Importantly, it also confirmed that our improved `diffSplice2` method outperforms the original, at no additional computing cost.

**Figure 2:**
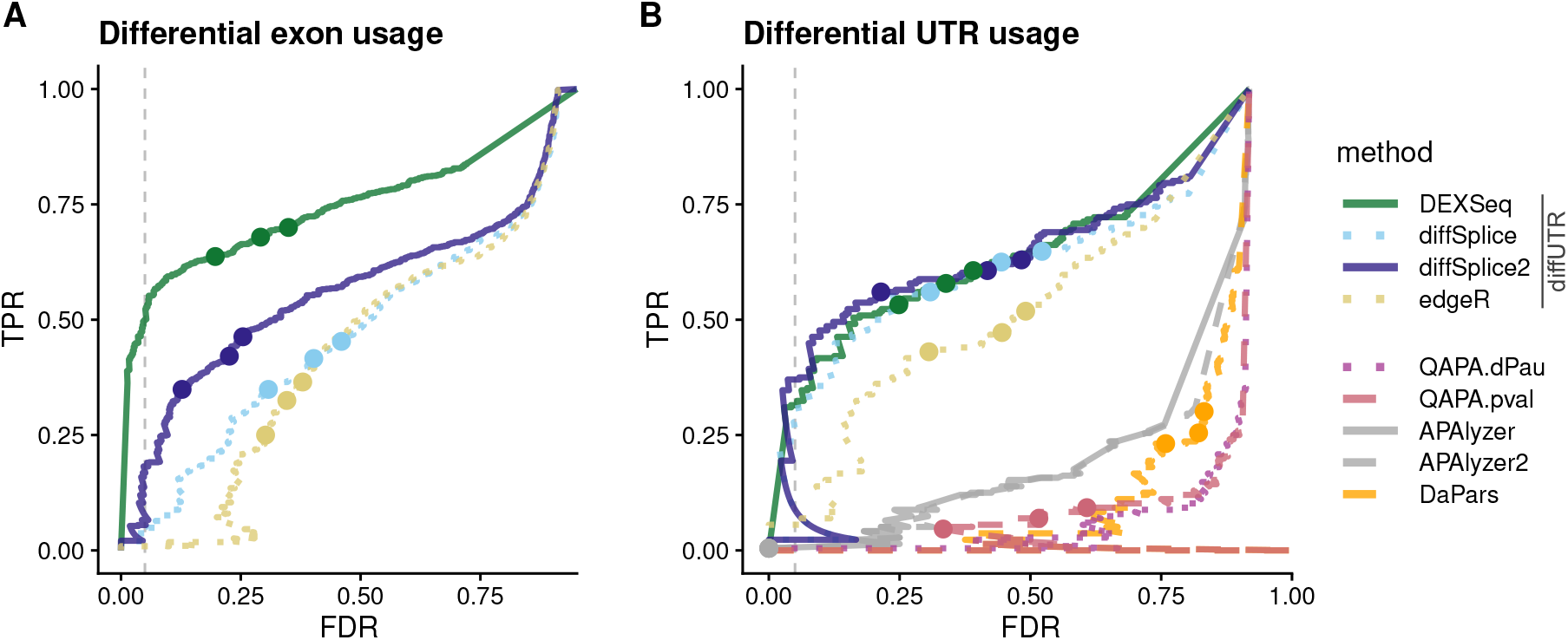
FDR and recall (TPR) on simulated data. **A:** In the classical DEU context. **B:** In the differential UTR usage context. The dashed line indicates a real False Discovery Rate (FDR) of 5%, and the dots indicate nominal FDRs of 10, 5 and 1%. *diffUTR* methods far outperform *QAPA* and *DaPars.* In both contexts, our modifications to diffSplice significantly improve its performance.

### Application to differential UTR usage and benchmark on a simulation

We next sought to evaluate the methods when applied for differential UTR analysis. For this purpose, APA sites are used to further segment and extend UTR bins, as illustrated in Figure 1B (see methods for the details). Given the absence of RNAseq data with a differential UTR usage ground truth, we simulated reads with known UTR differences from real data (see Simulated Data). We then ran the different *diffUTR* methods (as well as the unmodified diffSplice variant), and compared them to alternative methods. While *DaPars* and *APAlyzer* provide genelevel significance testing, *QAPA* does not, and our attempts to use its equivalence classes with standard transcript usage methods (see methods) gave very poor results. Therefore, for the purpose of comparison we tried two alternatives: simply ranked genes according to QAPA’s main output, i.e. the absolute difference in polyA site usage between conditions *(|ΔPAU*|), labeled in 2B as *QAPA.dPau,* or running t-tests on the log-transformed PAU values, labeled as *QAPA.qval.* Since *APAlyzer* produces different analyses for genes’ 3’ end and intronic APA usage, we used both the 3’ end results and a combination of the two (the latter shown as *APAlyzer2*). As Figure 2B shows, all *diffUTR* methods outperformed alternatives by far. On this test, our improved `diffSplice2` had comparable performance to *DEXSeq,* at a fraction of the computing costs.

### Differential UTR usage in real data

We next sought to test *diffUTR* in real data. First, since 3’ UTRs are known to generally lengthen during neuronal differentiation^[28,8]^, we expected to observe a skew towards positive fold changes of 3’ UTR bins when comparing RNAseq experiments from embryonic stem cells (ESC) and ESC- derived neurons. We therefore re-analyzed data from^[29]^ and observed clearly the expected skew among statistically-significant genes, especially for bins with a higher expression (Figure 3A).

**Figure 3:**
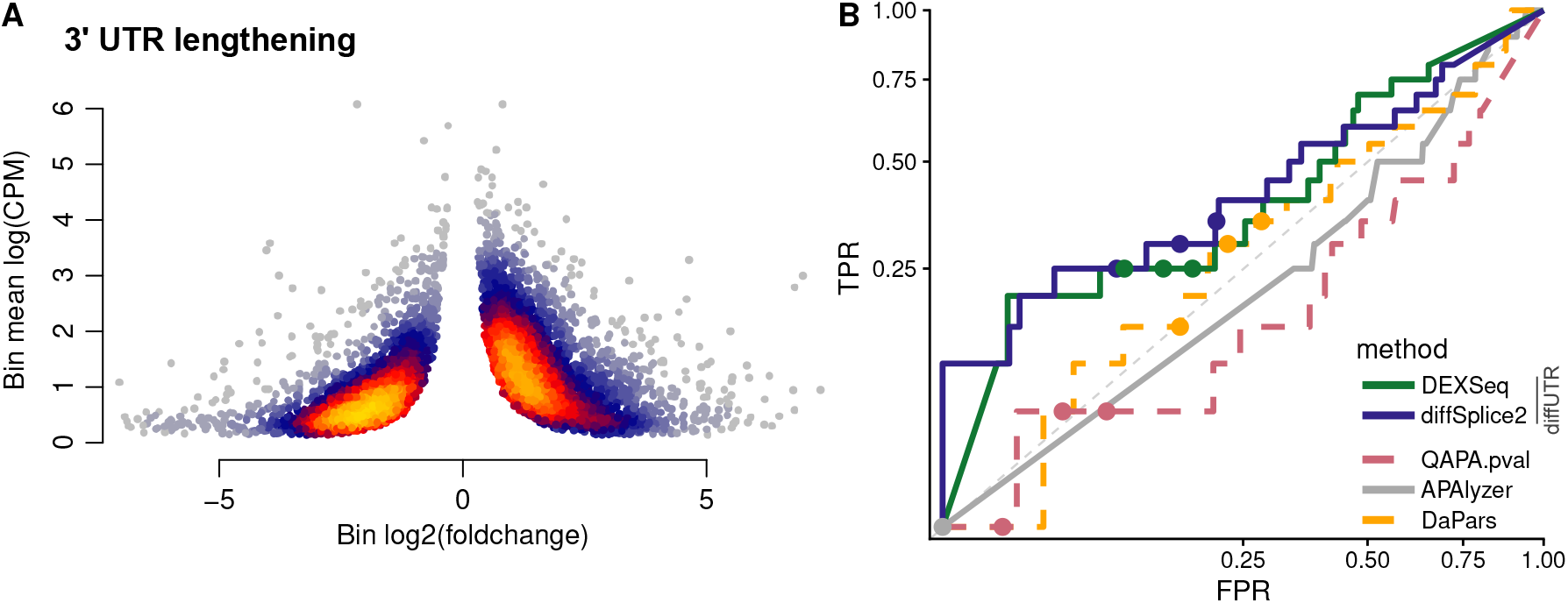
Differential UTR analysis on real data. **A:**. 3’ UTR lengthening during neuronal differentiation. Plotted are the UTR bins found statistically significant (bin- and gene-level FDR both i 0.1) by *diffUTR* (diffSplice2) when comparing in vitro differentiated neurons to mouse embryonic stem cells. The color indicates the point density. The clear skew towards a positive binlevel foldchange (indicative, in most cases, of a UTR lengthening), especially for bins with a higher mean count (CPM=counts per million reads sequenced). **B:** Receiver-operator characteristic (ROC) curves of differential UTR usage analysis on the LTP dataset, using 3’ sequencing to establish the ground truth. The axes are square-root-transformed to improve visibility, and only a subset of method variations are shown (see Supplementary Figure 1 for all variants).

We next found both 3’ sequencing and standard RNAseq data from samples of mouse hippocampal slices undergoing Forskolin-induced long-term potentiation^[30]^, which enabled us to use the 3’ sequencing data as a truth for analysis performed on the standard RNAseq data (Figure 3B and Supplementary Figure 1). In this case we represent the results through Receiver-operator characteristic (ROC) curves since the Precision-recall curves make the differences less visible due to the lower general power. Although power to detect UTR changes is necessarily low with respect to 3’ sequencing, we again observed that *diffUTR* methods clearly outperformed all alternative methods.

### Exploring differential exon/UTR usage results

*diffUTR* provides three main plot types to explore differential bin usage analyses, each with a number of variations. Figure 4 showcases them in the context of long-term potentiation of mouse hippocampal neurons^[30]^. plotTopGenes (Figure 4A) provides gene-level statistic plots (similar to a ‘volcano’ plot), which come in two variations. For standard DEU analysis, absolute bin-level coefficients are weighted by significance and averaged to produce gene-level estimates of effect sizes. For differential 3’ UTR usage, where bins are expected to have consistent directions (i.e. lengthening or shortening of the UTR) and where their size is expected to have a strong impact on biological function, the signed bin-level coefficients are weighted both by size and significance to produce gene-level estimates of effect sizes. By default, the size of the points reflects the relative expression of the genes, and the color the relative expression of the significant bins with respect to the gene.

**Figure 4:**
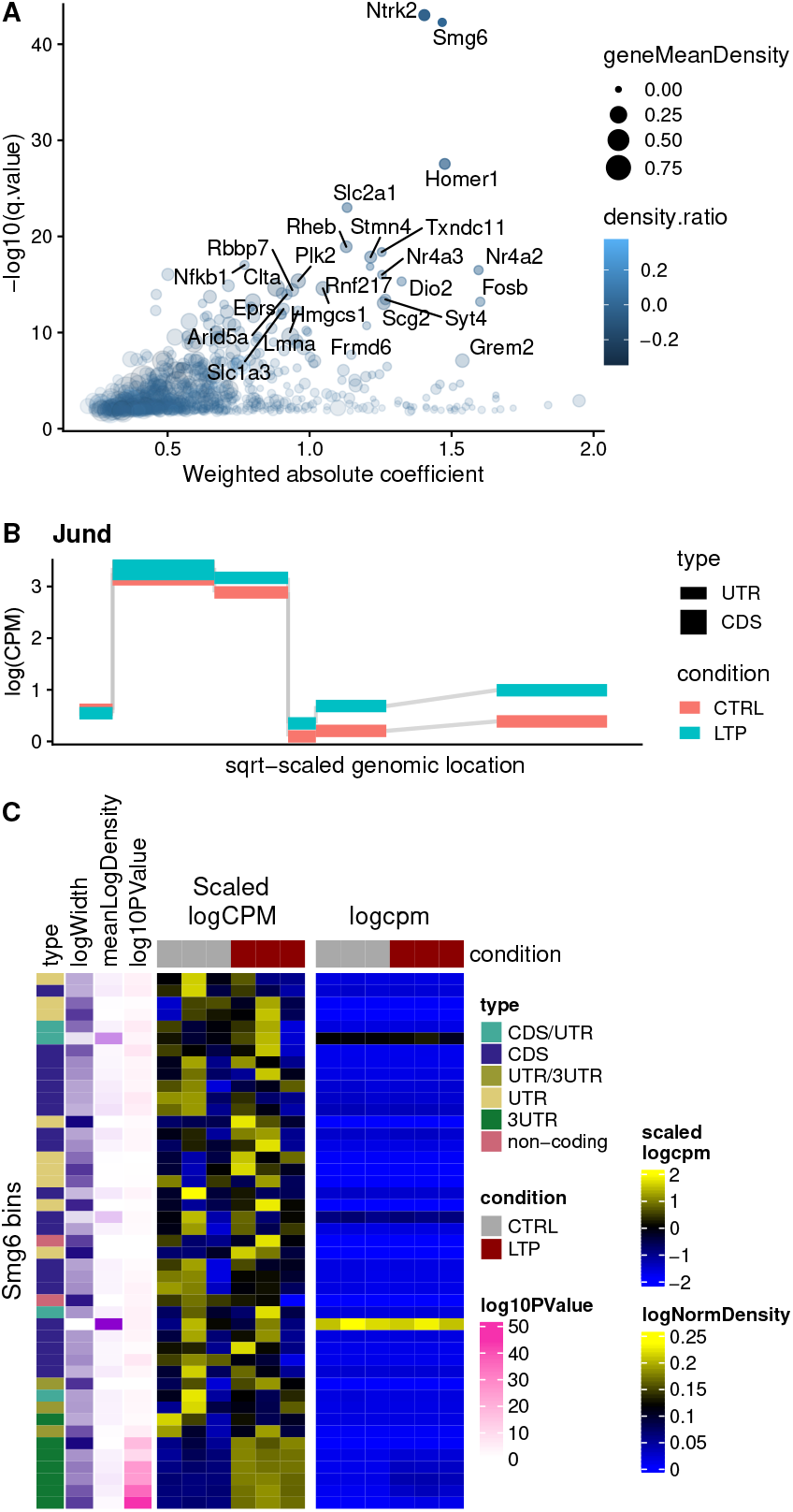
Plotting functions. **A:** `plotTopGenes` provides significance and effect size statistics aggregated at the gene level. **B:** `deuBinPlot` provides a more flexible version of the bin-level gene plots generated by common DEU packages. Shown here is the upregulation of Jund 3’ UTR upon LTP. **C:** `geneBinHeatmap` provides a compact, bin-per-sample heatmap representation of a gene.

`deuBinPlot` (Figure 4B) provides bin-level statistic plots for a given gene, similar to those produced by *DEXSeq* and *limma,* but offering more flexibility. They can be plotted as overall bin statistics, per condition, or per sample, and can display various types of values. Importantly, since all data and annotation are contained in the object, these can easily be included in the plots. Figure 4B shows a lengthening of the Jund 3’ UTR in the LTP group.

Finally, `geneBinHeatmap` (Figure 4C) provides a compact, bin-per-sample heatmap representation of a gene, allowing the simultaneous visualization of various information. We found these representations particularly useful to prioritize candidates from differential bin usage analyses. For example, many genes show differential usage of bins which are generally not included in most transcripts of that gene (low count density), and are therefore less likely to be relevant.

### Further variations tested

During implementation, we tested other changes to the method which were ultimately discarded as they did not improve performance, but which we here briefly report.

First, differential UTR analysis differs from typical differential exon usage analysis in that the vast majority of UTR bins are consecutively transcribed, meaning that changes in the usage of a bin should also be visible in downstream bins. We therefore reasoned that it would be beneficial to use this property to improve statistical analysis. We reasoned that connected bins with significant fold changes in the same direction could be unified and their p-values aggregated, and tested a rudimentary implementation using Fisher’s aggregation. However, this decreased accuracy and led to a worse FDR control (Supplementary Figure 2).

Second, most methods compare bin-level foldchanges to gene-level ones to identify bins behaving differently from the others, and we reasoned that, especially for genes with more UTR bins than CDS bins, including counts of 3’ UTR when calculating overall gene expression could under-estimate the gene expression and possibly mistake the UTR foldchange for the gene foldchange. We therefore tried a modification of *diffSplice* to only calculate the gene foldchange from coding sequence (CDS) bins and then compare it to the individual bins. Again, this approach proved unsuccessful (Supplementary Figure 3).

## Discussion

*diffUTR* streamlines DEU analysis and outperforms alternative methods in inferring UTR changes, which demonstrates the utility of harnessing powerful, well-established frameworks for new ends. It must be noted that the way in which the simulation was performed, i.e. elongating transcripts to the next polyA site(s), is similar to the way *diffUTR* disjoins the annotation into bins, which could cause a bias towards this method (as well as *QAPA* and *APAlyzer,* which also makes use of alternative polyA sites). However, this is unlikely to be the reason for the observed superiority of *diffUTR-based* methods given the considerable extent by which they outperformed alternatives, and the observation of similar results in real data.

Similar to DEU tools^[27]^, *diffUTR* fails to control the FDR correctly, and our attempts so far to improve this remained unsuccessful. We therefore recommend prudence with results close to the significance threshold. In addition, and in contrast to DEU where exons are subject to splicing in a potentially independent fashion, 3’ UTRs typically do not undergo splicing and therefore only differ in length between conditions. This means that the behavior of a UTR bin is dependent on that of upstream bins, a property which could be exploited to improve accuracy at the gene-level. However, our simple attempt to do so by combining p-values of consecutive bins did not have the desired outcome, pointing to the need of more research in this direction.

Further, the bin-based approach has the drawback of not pinpointing the exact UTR locations: it is limited to the bin resolution, and the bins themselves are limited by incomplete transcript and APA annotations. Additionally, because there is a significant drop off in read coverage at the end of transcripts, we have observed that it is often bins upstream of the actual UTR lengthen- ing/shortening event which give a statistically-significant signal rather than the one truly affected. This is why we have provided tools to enable the further inspection of events in a given gene.

Finally, the results of bin-based analyses are limited by the overlaps of transcripts from different genes, an issue on which differential transcript usage analysis approaches appear superior (e.g.^[31]^). However, transcript usage analysis tools are dependent on the completeness of the transcript annotation, while bin-based approaches are more open to the discovery of unannotated transcript variants, which is especially relevant for differential UTR usage. Here, we made the choice of including ambiguous bins, but flagging them as such, enabling users to interpret them with caution. While *DEXSeq* remains the tool of predilection for relative bin usage analyses, it scales very badly to larger sample sizes, and alternatives might be needed in some contexts. Our changes to *limma’s* original diffSplice method consistently result in more accurate predictions, making this new method the best compromise for bin-based approaches when *DEXSeq* is not applicable. More generally, it also shows that even with well-established approaches, there is still room for incremental, but non-negligible improvement.

## Methods

### 0.1 Data and code availability

The data objects and code used to produce the figures are available through the https://github.com/plger/diffUTR_paper repository. The *diffUTR* source code is available at https://github.com/ETHZ-INS/diffUTR.

### 0.2 RNAseq data processing

For the evaluation of `diffSplice2` in a standard DEU case, we used bin count data obtained from the authors of the original DEU benchmark^[27]^. For other datasets, reads were downloaded from the SRA, aligned to the GRCm38.p6 genome using STAR 2.7.3a with default parameters and the GENCODE M25 annotation as guide. The same gene annotation was used as input for bin creation.

### 0.3 diffUTR

*diffUTR* is implemented as a Bioconductor package making use of the extensive libraries avail-able, especially the *GenomicRanges* package^[32]^ and the different DEU methods (see Differential analysis).

#### 0.3.1 Preparing bins

Exons are extracted from the genome annotation and flattened into non-overlapping bins (Figure 1B). In other words, the exon annotation is fragmented into the widest ranges where the set of overlapping features is the same. Bins that do not overlap with coding sequences (CDS) and belong to a protein coding transcript are labeled as UTR and the rest as CDS. When APA sites are also provided as input (for the purpose of this article, polyAsite v2.0 sites were used), bins are further segmented and/or extended. For this the closest upstream CDS or UTR is found for every poly(A) site and the UTR is defined from this boundary to the polyA site and assigned to the corresponding gene and transcript (Figure 1B). If the newly defined UTRs exceeds a predefined length specified by `maxUTRbinSize` (default is 15000bp), it is ignored as unlikely to be a real UTR. Moreover, if the start of a gene is the closest upstream sequence before any UTR or CDS the newly defined UTR is ignored to avoid assignment problems. In order to later differentiate between regions that are 3’ or 5’ UTRs, regions that are downstream of the last CDS of a given transcript were labeled as 3’ UTR. The label ‘non-coding’ is assigned to all bins that have no protein coding transcript overlapping it.

If a bin originates from regions belonging to different genes, the bin is duplicated and assigned once to each gene, so that each gene contains the same fragment once. Alternatively, the `genewise` argument can be used so that only exons belonging to the same gene are considered when flattening.

#### 0.3.2 Quantification

For quantification, `countFeatures()` uses the `featureCounts()` function from the *Rsubread* `package`^[33]^ to count previously mapped reads overlapping each bin. By default every read is assigned once to every bin it overlaps with and can therefore be counted multiple times, which is needed because many bins are shorter than the read length. Alternative counting methods, such as `summarizeOverlaps()` from the *GenomicAlignments* package^[32]^ performed considerably worse in the simulation. The function returns a `RangedSummarizedExperiment` object^[26]^, containing the read counts as well as the bin annotation.

#### 0.3.3 Differential analysis

Three wrappers implement corresponding DEU methods on the `RangedSummarizedExperiment` object previously generated, returning results as further standardized annotation within the object. For differential UTR analysis, gene-level results are obtained by filtering the bin-level results for those assigned to the type UTR and/or 3’ UTR, and setting all other p-values to 1 before aggregation.

**diffSpliceDGE.wrapper()** This is a wrapper around *edgeR*’s DEU method based on fitting a negative binomial generalized linear model^[23]^. In a first step the bins are filtered to decide which have a large enough read count to be kept for the statistical analysis (filterByExpr()), the library sizes are normalized (`calcNormFactors()`) and the dispersion is estimated (`estimateDisp()`). After this the model is fitted (`glmFit()`). If the option `QLF = TRUE` (default), an extended model is fitted, using quasi-likelihood methods to account for gene specific variability (`glmQLFit()`). In the last step bin fold changes are tested to be different from overall gene fold changes, using a likelihood ratio test or a quasi-likelihood F-Test depending on the `QLF` option chosen (`diffSpliceDGE()`). The gene level p-values are obtained by the Simes’ method^í34]^.

**DEXseq.wrapper()** In this method the standard *DEXseq* differential exon usage pipeline^[22]^ is implemented. It is similarly to edgeR based on fitting a negative binomial model but instead of comparing fold change differences between bins and genes, *DEXseq* compares a full model containing a term corresponding to the change in exon usage between conditions to a reduced model without this term. The two fits are compared using a χ^2^ likelihood-ratio test. The libraries are normalized (`estimateSizeFactor()`), the dispersion is estimated (`estimateDispersion()` and the models are fitted (`testForDEU()`). In a last step the fold changes between the bins are estimated (`estimateExonFoldChanges()`). To obtain gene level results the function `perGeneQValue()` is used, which is based on the Šidák method^[35]^.

**diffSplice.wrapper() and diffSplice2** This method implements the differential exon usage pipeline of *limma* for RNA-seq data^[25]^. The pre-processing is identical to `diffSpliceDGE.wrapper()`, then the precision weights are estimated with (`limma::voom()`) and the linear models are fitted (`limma::lmFit()`). In the last step, bin fold changes are tested to be different from overall gene fold changes, using a moderated t-test (`diffSplice()` or, by default, `diffSplice2()` – see below). The gene level p-values are obtained by the Simes’ method^134]^.

The `diffUTR::diffSplice2` function provides an improved version of *limma’s* original `diffSplice` method. `diffSplice` works on the bin-wise coefficient of the linear model which corresponds to the log2 fold changes between conditions. It compares the log2(fold change) 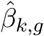 of a bin *k* belonging to gene *g*, to a weighted average of log2(fold change) of all the other bins of the same gene combined 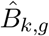 (the subscript *g* will be henceforth omitted for ease of reading). The weighted average of all the other bins in the same gene is calculated by

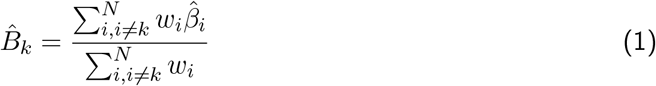

where 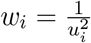 and *u_i_* refers to the diagonal elements of the unscaled covariance matrix (*X^T^VX*)^-1^. *X* is the design matrix and *V* corresponds to the weight matrix estimated by `voom`. The difference of log2 fold changes, which is also the coefficient returned by `diffSplice()` is then calculated by 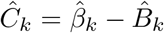. Instead of calculating the t-statistic with 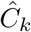, this value is scaled again in the original code:

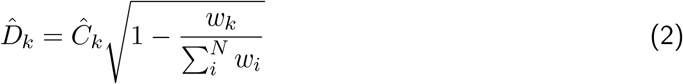

and the t-statistic is calculated as:

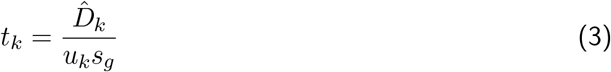

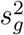 refers to the posterior residual variance of gene *g*, which is calculated by averaging the sample values of the residual variances of all the bins in the gene, and then squeezing these residual variances of all genes using empirical Bayes method. This assumes that the residual variance is constant across all bins of the same gene.

In `diffSplice2()`, we applied three changes to the above method. First, the residual variances are not assumed to be constant across all bins of the same gene. This results in the sample values of the residual variances of every bin now being squeezed using empirical Bayes method, resulting in posterior variances 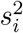 for every individual bin *i*. Second, the weights *w_i_*, used to calculate 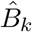, now incorporate the individual variances by 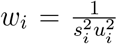. Third, the 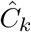 value is directly used to calculate the *t*-statistic, which after all these changes now corresponds to

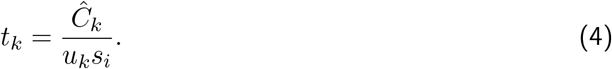

### 0.4 Simulated Data

The simulation was done using the *Polyester* R package^[36]^ using parameters obtained from the control samples of mouse hippocampus RNAseq^[30]^. Using *salmon*^[17]^ with a decoy-aware transcriptome index for the mm10 genome from^[37]^, the abundances for each transcript were first estimated to learn parameters for the simulation. 1000 transcripts from different genes were randomly chosen. The last exon of all these transcripts was lengthened to the next, second next or third next downstream APA site annotated in the polyAsite database^[15]^. Duplicates of these transcripts were generated, which had less or no lengthening of their last exon, generating pairs of transcripts with different UTR lengths. For each transcript pair, one transcript was up and the other one down regulated by the same sampled fold change between 1.3 and 5. To make it more realistic, fold changes were also assigned to 300 genes from the set with differential UTR, and 300 genes that did not have differences in UTR usage. Reads were then generated for two conditions with three replicates each using the `simulate_experiment()` function with the options `paired = FALSE, error_model = “illumina5”, bias = “cdnaf”` and `strand_specific = TRUE`. The simulated reads are available on figshare at https://dx.doi.org/10.6084/m9.figshare.13726143.

### 0.5 3’-seq analysis

To establish a set of true relative differences in UTR usage from the 3’ sequencing data^í30]^, we downloaded the authors’ counts per cluster from the Gene Expression Omnibus (file `GSE84643_3READS_count_table.txt.gz`). We used the 3h treatment because we observed it to have the strongest signal, and excluded one sample (A6) that appeared like a strong outlier based on PCA and MDS plots. We kept only clusters with at least 50 reads in at least 2 samples, and used *DEXSeq* to fit a negative binomial on each gene and estimate the significance of the `cluster:condition term`. We considered as true positives genes with a gene-level and bin-level q-value ≤ 0.1, and true negatives genes with a gene-level q-value ≥ 0.8. Genes for which all tested methods produced a p-value of 1 or NA (i.e. genes filtered out as too lowly expressed in the standard RNAseq) were excluded for the benchmark.

### 0.6 Comparisons with alternatives

For the comparison of methods, all functions were used with their default parameters and run according to their manual. As *QAPA* and *DaPars* do not provide means to aggregate the results to the gene level this was implemented separately. For *DaPars* the p-values were aggregated to the gene level by using Simes’ method^[34]^ for comparability with *diffUTR.* Aggregation by taking the minimum p-value of all the transcripts in a gene produced extremely similar results. For *QAPA* |ΔPAU| was calculated and aggregated to a gene level by taking the maximum from all transcripts of a gene and the genes were ranked by this value. Alternatively, we also tested applying a t-test on the log-transformed *PAU* values (log-transforming had a negligible effect), followed by Simes’ gene-level aggregation. Attempts to complement *QAPA* with p-values estimated from established statistical tests working with its equivalence classes, such as BANDITS^[31]^, did not improve the results and were therefore discarded so as not to distort the original method. Finally, for *APAlyzer2* we combined the 3’ UTR and intronic APA analyses by using the minimum of the two p-values. See the https://github.com/plger/diffUTR_paper repository for details.

We used the following software versions for comparisons: *Polyester* 1.24.0, *DEXSeq* 1.34.0, *edgeR* 3.30.0, *limma* 3.44.0, *DaPars* 0.9.1, *APAlyzer* 1.5.5. For QAPA, we used *salmon* 1.3.0 with validateMappings.

## Author’s contributions

SG developed the bin preparation and the diffSplice modification, and ran most of the analyses. PLG and SG wrote the package and paper. PLG and GS supervised the project.

## Acknowledgements

SG performed this research as part of his bachelor thesis in the Interdisciplinary Sciences program at ETH. PLG’s position is co-funded by Prof. Mark Robinson (Institute of Molecular Life Sciences, University of Zurich) and Professors Gerhard Schratt, Johannes Bohacek and Isabelle Mansuy (Institute of Neuroscience, ETH Zurich). GS is supported by grants from the SNF (SNF_179651, SNF_189486) and the ETH (ETH-24 18-2 (NeuroSno)). We thank the Robinson group (UZH) for feedback.

